# The lncRNA *EPB41L4A-AS1* regulates gene expression in the nucleus and exerts cell type-dependent effects on cell cycle progression

**DOI:** 10.1101/2021.02.10.430566

**Authors:** Helle Samdal, Siv Anita Hegre, Konika Chawla, Nina-Beate Liabakk, Per Arne Aas, Bjørnar Sporsheim, Pål Sætrom

## Abstract

The long non-coding RNA (lncRNA) *EPB41L4A-AS1* is aberrantly expressed in various cancers and has been reported to be involved in metabolic reprogramming and as a repressor of the Warburg effect. Although the biological relevance of *EPB41L4A-AS1* is evident, its functional role seems to vary depending on cell type and state of disease. By combining RNA sequencing and ChIP sequencing of cell cycle synchronized HaCaT cells we previously identified *EPB41L4A-AS1* to be one of 59 lncRNAs with potential cell cycle functions. Here, we demonstrate that *EPB41L4A-AS1* exists as bright foci and regulates gene expression in the nucleus in both *cis* and *trans*. Specifically, we find that *EPB41L4A-AS1* positively regulates its sense overlapping gene *EPB41L4A* and influences expression of hundreds of other genes, including genes involved in cell proliferation. Finally, we show that *EPB41L4A-AS1* affects cell cycle phase distribution, though these effects vary between cell types.

## Introduction

According to the Encyclopedia of DNA elements (ENCODE) project (GENCODE v35) there are 17957 annotated lncRNA genes giving rise to 46977 different transcripts. LncRNAs are characterized as transcripts more than 200 nucleotides long with none or little coding potential. Most of them are transcribed by RNA polymerase II (Pol II), poly-adenylated, 5’-capped, and spliced. Functional evaluation of lncRNAs demonstrate that they are involved in gene regulation at the transcriptional, epigenetic, and translational level, where they can interact with DNA, RNA, and proteins to exert their function. LncRNAs can broadly be classified based on whether they regulate gene expression around their transcription site in *cis*, or if they leave their site of transcription and exert their function elsewhere in *trans* [1]. Several lncRNAs are involved in the development and pathogenesis of diseases, including different types of cancer, degenerative, autoimmune, and cardiovascular diseases. Although their biological relevance is undisputable, only a small part of the annotated lncRNAs have been functionally evaluated.

As RNA is the functional product of lncRNAs, their subcellular localization provides information about their potential biological role. Nuclear localized lncRNAs have possible roles connected to chromatin organization, transcriptional regulation, RNA processing, or nuclear domains. Meanwhile, cytoplasmic lncRNAs can modulate gene regulation by acting as decoys for microRNAs (miRNAs) and proteins, or affect signal transduction by interfering with protein post-translational modifications [2]. LncRNAs are more cell type-specific than mRNAs, and both their subcellular localization and biological role can vary according to cell/tissue and stage of development [2]. This characteristic increases the complexity and adds to the challenge in determining the biological role of lncRNAs. For instance, Metastasis Associated Lung Adenocarcinoma Transcript 1 (*MALAT1*) is one of the best-studied lncRNAs, and it is implicated in several diseases such as diabetes, atherosclerosis, and cancer. *MALAT1* is involved in different cellular processes including differentiation, tumor progression, hypoxia, inflammation, and stress. There are accumulating studies demonstrating an oncogenic role of *MALAT1* [3], and interestingly, a few studies reporting a tumor-suppressor role [4–7].

*EPB41L4A-AS1* is identified as dysregulated in several gene expression studies of different diseases, suggesting a possible role as biomarker and a therapeutic target [8, 9]. There are a few functional studies investigating the biological role and molecular mechanisms of *EPB41L4A-AS1* [9–12] (Hegre and Samdal *et.al*. unpublished). Two functional studies report that *EPB41L4A-AS1* is involved in the regulation of glycolysis and glutaminolysis, and acts as a suppressor of the Warburg effect in the placental villus and in cancer cells [11, 12]. Another functional study demonstrates that mir-146a inhibits the expression of *EPB41L4A-AS1*, and overexpression of *EPB41L4A-AS1* affects both proliferation and cell cycle phase distribution in bone marrow-derived mesenchymal stem (BMSC) cells [10].

In a previous study from our group we combined RNA sequencing (RNA-seq) data with ChIP-sequencing (ChIP-seq) signal from cell cycle synchronized HaCaT cells where we identified 94 lncRNAs with a cyclic expression profile. Of these, 59 had a positive correlation to Pol II or histone 3 lysine 4 trimethylation (H3K4me3) signals, consistent with being actively transcribed at specific phases of the cell cycle (Hegre and Samdal *et.al.* unpublished). Among these lncRNAs, we selected *EPB41L4A-AS1* for further functional evaluation. We observed a decreased proliferation in HaCaT, A549, and DLD1 cells in response to siRNA-mediated downregulation of *EPB41L4A-AS1*. In addition, siRNA-mediated knockdown of *EPB41L4A-AS1* led to a reduction of cells present in the G1 phase and an enrichment of cells present in the G2/M phase of the cell cycle in HaCaT cells, indicating a possible role in the G2/M progression.

Here, we used RNA FISH and determined that the subcellular localization of *EPB41L4A-AS1* is both nuclear and extranuclear. We used CRISPR interference and CRISPR activation (CRISPRi and CRISPRa, respectively) to modulate the expression of *EPB41L4A-AS1* at the transcriptional level, followed by viability and cell cycle assays. In line with our previous study, we find that CRISPR-mediated downregulation affects cell cycle phase distribution. The effect on phase distribution was different between HaCaT and A549 cells, suggesting that *EPB41L4A-AS1* has cell type-specific roles. Moreover, *EPB41L4A-AS1* has *cis*-regulatory abilities, and is a positive regulator of its antisense mRNA neighbor *EPB41L4A.* CRISPR-mediated upregulation of *EPB41L4A-AS1* resulted in the opposite effect on phase distribution and *EPB41L4A* expression level. Finally, we used RNA-seq to provide mechanistic insight into the functional role of *EPB41L4A-AS1.* The analysis suggests that *EPB41L4A-AS1* affects the expression of multiple genes in HaCaT and A549 cells, and gene ontology (GO) terms include several growth promoting signaling pathways.

## Material and Methods

### Cell culture

Cell lines were obtained from the American Type Culture Collection (ATCC) and cultivated in a humidified incubator in 5% CO_2_ at 37°C. HaCaT, A549, and Hek293T cells were all cultured in Dulbecco’s modified Eagle’s medium (DMEM) (Sigma-Aldrich, D6429) supplemented with 10% fetal bovine serum (FBS) (Sigma-Aldrich, F7524) and 2 mM L-Glutamine (Sigma-Aldrich, G7513). For XTT viability assays we used DMEM without phenol red (Thermo Fisher Scientific, 21063029). All cell lines were sub-cultivated at about 70 % confluence at least twice a week.

### Single molecule RNA fluorescence in situ hybridization (RNA FISH)

We used the software available through Stellaris Probedesigner to design the oligonucleotide set for RNA FISH. For the *EPB41L4A-AS1* we used masking 3, length 20, spacing 2 which resulted in 36 oligonucleotides. Stellaris™ probesets (Biosearch Technologies) were conjugated to a Quasar670 dye in the 3′ end. Hybridization and staining were performed as prescribed in the Stellaris protocol for adherent cells. GAPDH was used as a predesigned control. The different probe sequences are listed in Supplementary Table 1.

### Imaging

To image the cells, we used a Zeiss Laser TIRF 3 fluorescence microscope (Zeiss), equipped with a α-Plan-Apochromat 100x/1.46 oil-immersion objective. We used a Zeiss 81 HE DAPI/FITC/Rh/Cy5 filter, DAPI was excited by LED-module 365 nm (Zeiss Colibri) and Quasar670 was excited by 644 nm wavelength laser. The images were acquired by either a Hamamatsu EMCCD EMX2 or a Hamamatsu ORCA-Fusion camera at 16 bit and at a voxel size of 100 × 100 × 220 nm^3 (EMCCD) or 129 × 129 × 220 nm size (ORCA-Fusion). We used SVI Huygenes Professional (version 18.10) for image deconvolution and image analysis was done using Fiji (version 1.52t) [13]. Presented images are maximum intensity projections of 34 Z-stack slices (7.26 μm) of the cell.

### Plasmids and cloning

We followed the Zhang labs protocol (https://media.addgene.org/data/plasmids/52/52961/52961-attachment_B3xTwla0bkYD.pdf) for guide RNA (gRNA) design and cloning of the gRNA between the two BsmBI restriction sites. Lenti-sgRNA puro was a gift from Brett Stringer (Addgene plasmid # 104990). The hU6-F (5’-GAGGGCCTATTTCCCATGATT-3’) was used to sequence the RNA to validate gRNA insert. For CRISPRa we used pXPR_120 [14] with multiple activating domains, VP64, P65 and Rta, and for CRISPRi we used pHR-SFFV-dCas9-BFP-KRAB [15], a gift from John Doench & David Root (Addgene plasmid # 96917) and Stanley Qi & Jonathan Weissman (Addgene plasmid # 46911), respectively.

### Lentiviral production

We plated Hek293T cells 24 hours before transfection with Lenti-sgRNA puro containing the different gRNAs or pHR-SFFV-dCas9-BFP-KRAB or pXPR_120 together with the packaging plasmids psPAX.2 and pMD2.G. Lipofectamine 2000 (Invitrogen™, 11668019) was used as a transfection reagent. We replaced the culture medium 8 hours after transfection to decrease toxicity. We collected the viral supernatant 72 hours after transfection, centrifuged at 1800 g for 5 min at 4°C, and filtered it through a 45 μM filter. The virus was stored at −80℃.

### Generating stable dCas9 expressing cell lines

Both HaCaT and A549 cells were transfected by adding lentiviral titer of pXPR_120 or pHR-SFFV-dCas9-BFP-KRAB together with 8 μg/ml polybrene in DMEM. We replaced the medium 48 hours after transduction. After transduction, 10 μg/ml Blasticidin (Invivogen) was used to select for cells that had incorporated pXPR_120 into their genome. We changed the medium after three days, while selection was continued for about seven days. To keep the selection pressure, we sub-cultivated the pXPR_120 cells with 5 μg/ml Blasticidin. We used a FACS Aria II cell sorter (BD Bioscience) to select cells transduced with pHR-SFFV-dCas9-BFP-KRAB.

### Guide RNA (gRNA) design

We used the E-CRISPR online tool (http://www.e-crisp.org/E-CRISP) for gRNA design [16]. The sequence targeted −50 to + 200 base pairs (bp) relative to the transcriptional start site (TSS) of *EPB41L4A-AS1* (hg38_dna range=chr5:112160660-112160910). The exact location of TSS was determined using Fantom5 web resource (ZENBU 3.0.1) [17, 18]. To avoid off-target effects, we used Basic local alignment Search Tool to discard any sequences that were located within annotated genes other than target. The sequences of different gRNAs are listed in Supplementary Table S2.

### gRNA transductions

The cells were transduced with target-specific lentiviral gRNAs or a non-target gRNA together with 8 μg/ml polybrene approximately 20 hours after seeding at about 50% confluence. The multiplicity of infection (MOI) was 2. We added 2 μg/ml puromycin (Invivogen, ant-pr-1) to the growth media 24 hours after transduction to select for resistant cells containing the gRNA and harvested the cells 72 hours after selection was added. We included gRNA specific for *MALAT1* and *SCLA41* as positive controls for CRISPRi and CRISPRa, respectively.

### Short interfering (siRNA)-mediated knockdown

All cells were transfected with siRNAs for a 20 nM final concentration using Lipofectamine RNAimax (Invitrogen™, 13778030) when plated, according to the manufacturer’s protocol. Cells were harvested after 48 and/or 72 hours after transfection at about 70% confluence. MISSION® siRNA Universal Negative Control #1 (Sigma, SIC001) was used as control for siRNAs. The producers and sequences of siRNAs are listed in Supplementary Table S3.

### Quantitative Real-Time PCR (RT-qPCR)

The *mir*Vana miRNA Isolation Kit (Invitrogen™, AM1560) was used to isolate total RNA according to manufacturer’s protocol, followed by DNase treatment with TURBO DNA-*free*™ (Invitrogen™, AM1907). RNA concentration and quality were measured on a NanoDrop ND-1000 UV-Vis Spectrophotometer. RNA was converted to cDNA under standard conditions with random hexamer primers using TaqMan™ Reverse Transcription reagents (Invitrogen™, N8080234). Quantitative RT PCR reactions were prepared with Syber select master mix (Applied Biosystems, 4472919). The sequences of the different primers are listed in Supplementary Table S4. Relative expression levels were calculated using the comparative C_T_ method (2^−ΔΔCT^ method, [19]), and expression data were normalized to GAPDH.

### Viability assay

We performed TACS XTT Cell viability Assay (R&D Systems™, 4891025K) to investigate how down- and upregulation of *EPB41L4A-AS1* affected the viability, measured as the level of metabolic activity, in HaCaT cells. We transduced HaCaT cells with a non-target gRNA or an *EPB41L4A-AS1-*specific gRNA and harvested after selection with puromycin. Cells were then seeded in triplicates for each condition in a 96-well tray 96 hours after transduction. We measured absorbance 24, 48, and 72 hours after seeding, 4 hours after XTT was added, following the manufacturer’s protocol. We also performed cell counting using Moxi z mini automated cell counter (ORFLO Technologies) to investigate how CRISPRi-mediated downregulation of *EPB41L4A-AS1* affected proliferation in HaCaT cells. Cells were seeded in triplicates for each condition in a 24-well tray and counted 48 and 72 hours after transfection. Each well was washed twice with preheated PBS and trypsinated for 8-10 min before the cells were resuspended in preheated growth medium and counted.

### Cell cycle and fluorescence-activated cell sorting (FACS) analysis

For FACS analysis of the cell cycle phase distribution we harvested the cells 96 hours after gRNA transduction, by washing twice with preheated PBS and trypsinating for 8-10 min. Then we collected the cells using ice-cold PBS supplemented with 3% FBS, centrifuged the cells at 4°C for 5 min, and removed the supernatant. Cells were resuspended in 100 μl cold PBS, fixed in 1 ml ice-cold 100% methanol and stored at 4°C until DNA measurement. The cells were washed with cold PBS and incubated with 200 μl of DNase-free RNAse A in PBS (100 μg/ml) for 30 min at 37°C before DNA staining with 200 μl of Propidium Iodide (PI, Sigma) (50 μg/ml) at 37°C for 30 min. We performed cell cycle analyses by using a BD FACS Canto flow cytometer (BD Biosciences). The excitation maximum of PI is 535 nm and the emission maximum 617 nm. PI stained cells were excited with the blue laser (488 nm), and the PI fluorescence was detected in the PE channel (578 nm). We used FlowJo software to quantify the cells in each phase, and the percentage of cells that was assigned to G1, S, and G2/M phases was calculated.

### Total RNA sequencing (RNA-seq)

We used the *mir*Vana miRNA Isolation Kit (ThermoFisher Scientific, AM1560) to isolate total RNA, according to the manufacturer’s protocol. RNA concentration was measured on a Qubit (Thermo fisher), whereas integrity and stability of the RNA samples were assessed by using an Agilent 2100 Bioanalyzer (Agilent Technologies). The ribosomal RNA (rRNA) was removed using RiboCop rRNA Depletion Kit for Human/Mouse/Rat V2 (Lexogen, 037), and total RNA-seq libraries were prepared using CORALL (Lexogen, 095), according to the manufacturer’s instructions. The samples were sequenced using the Illumina NextSeq 500 Sequencing System.

### RNA-seq data analysis

We filtered and trimmed FASTQ files (fastp v0.20.0) and generated transcript counts using quasi alignment (Salmon v1.3.0) to the transcriptome reference sequence (Ensembl, GRCh38 release 92). The transcript sequences were imported into the R statistical software and aggregated to gene counts using the tximport (v1.14.0) bioconductor package. The genes with expression of less than 1 count per million in at least 50% of the samples were filtered out to compare gene expression between samples. We transformed count matrices by using the bioconductor package limma [20] combined with voom transformation [21]. Differentially expressed genes were identified using the toptable function of limma and we used the Benjamini-Hochberg method to correct for multiple testing. Genes with p-values < 0.05 were considered significantly differentially expressed.

## Results

### *EPB41L4A-AS1* is localized both in and outside the nucleus

The relative concentration index (RCI) from RNA-seq analyses of cytoplasmic and nuclear fractions in multiple cell lines suggest that *EPB41L4A-AS1* is mainly localized in the cytoplasm (Figure 1A; average RCI of 1.12). To address the question of *EPB41L4A-AS1*’s cellular localization in detail, we imaged HaCaT cells following RNA FISH against *EPB41L4A-AS1*. The data showed that in HaCaT cells, *EPB41L4A-AS1* appears as bright foci and low-intensity spots both in and outside the nucleus (Figure 1B; Supplementary Figure S1). Cross-sections of the nucleus confirmed that some bright foci co-localized with the DNA (Figure 1C).

**Figure 1:**
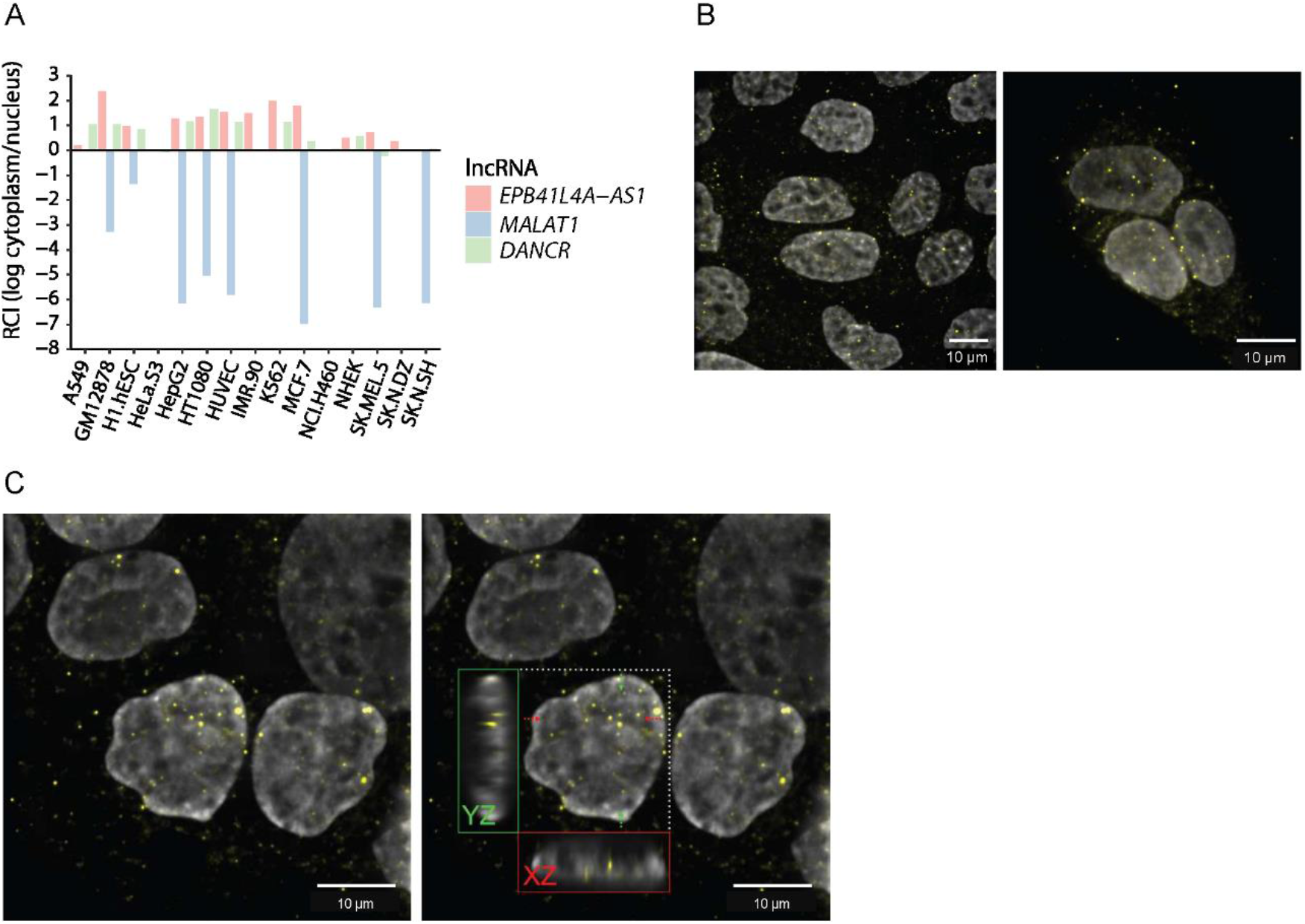
*EPB41L4A-AS1* has a nuclear and extranuclear localization. (A) The subcellular localization of *EPB41L4A-AS1*, *MALAT1,* and *DANCR.* Data are from the lncAtlas (https://lncatlas.crg.eu/) and display the subcellular localization based on the RCI of RNA between cytoplasm and nucleus. Values > 0 indicate cytoplasmic enrichment. (B) RNA FISH showing the localization of EPB41L4A-AS1 (yellow) in fixed HaCaT cells stained with DAPI (grey). Images are presented as maximum intensity projections of 34 Z-stack slices (7.26 μm) of the cell. (C) Cross section of the nucleus of fixed HaCaT cells shows *EPB41L4A-AS1* (yellow) localized with DNA (DAPI, grey). Images are presented as maximum intensity projections of 34 Z-stack slices (7.26 μm) of the cell. The red and green boxes show the signals from the cell planes marked by the red and green arrows projected onto the XZ and YZ planes, respectively.

### Generation of stable dCAS9 expressing cell lines

HaCaT is a spontaneously transformed human keratinocyte cell line and A549 is an adenocarcinoma human alveolar basal epithelial cell line. We successfully modified both cell lines to express catalytically inactive CAS9, a nuclease dead CAS9 (dCAS9), fused with either the Krüppel-associated box (KRAB) repressing domain or multiple activating domains to use as tools for transcriptional inhibition (CRISPRi) and activation (CRISPRa), respectively. The same two gRNAs were used for both up- and downregulation of *EPB41L4A-AS1*, although for A549, only gRNA 1 was able to downregulate the expression of *EPB41L4A-AS1* (Figure 2A).

**Figure 2.**
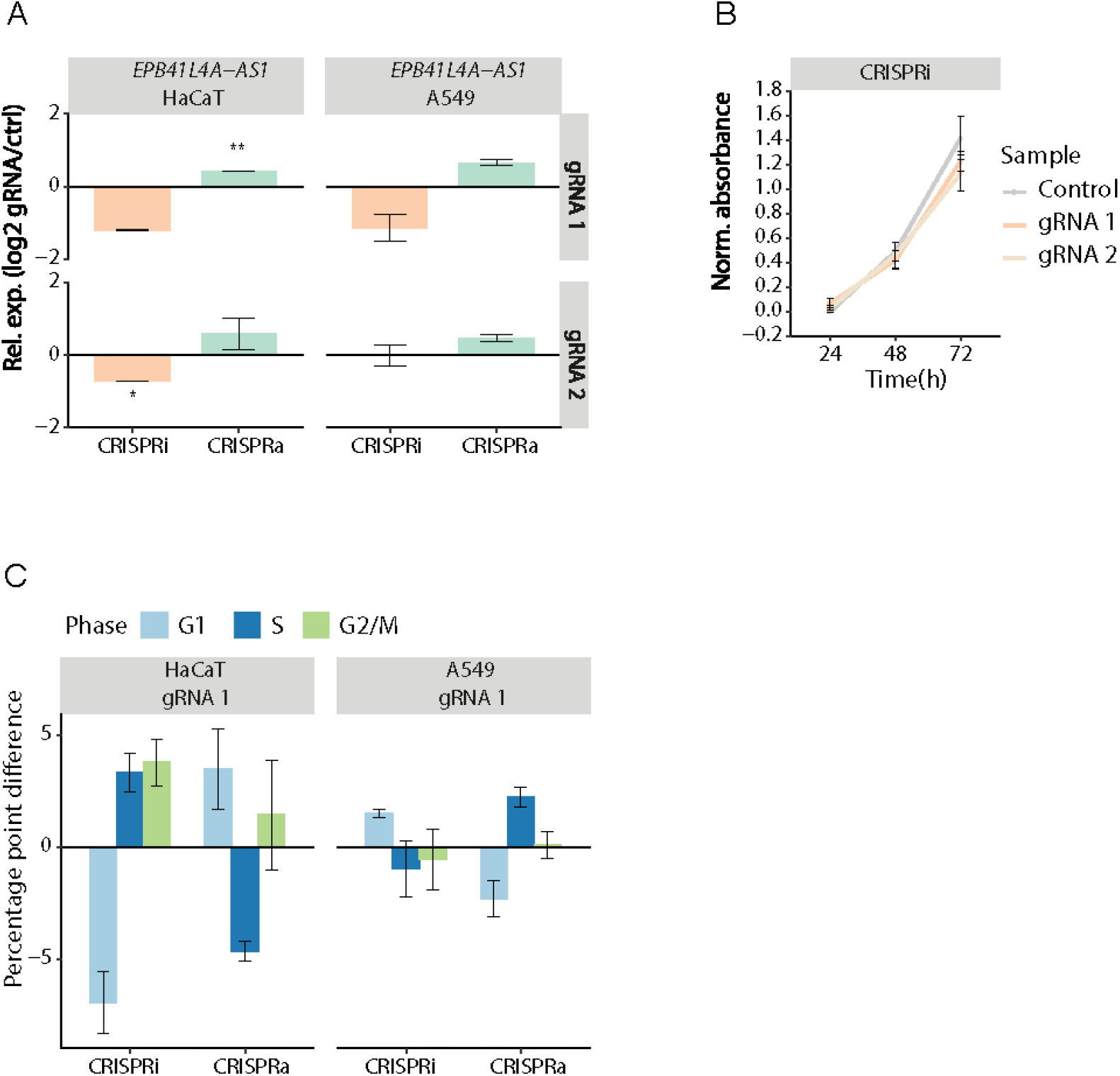
*EPB41L4A-AS1* affects cell cycle distribution. (A) The relative expression level of *EPB41L4A-AS1* as measured by RT-qPCR in response to CRISPRi/a of *EPB41L4A-AS1* in HaCaT and A549 cells. Data are presented as fold change expressions of cells transduced with gRNA 1 and gRNA 2 relative to control gRNA. Bars and error bars are mean and standard error of mean (SEM) of two independent replicates. Significant differences were determined by Student’s *t*-test, (unpaired, two-tailed) assuming equal variances (\**p* ≤ 0.05, \*\**p* ≤ 0.01). (B) Effect of CRISPRi of *EPB41L4A-AS1* on the metabolic activity in HaCaT cells, as measured by XTT. Data are presented as normalized absorbance (A_465 nm_ – A_630 nm_). Bars and error bars are mean and SEM of three independent replicates. *P*-values were determined by Student’s *t*-test, (paired, two-tailed) assuming equal variances (gRNA 1 *p* = 0.111 and *p* = 0.474, gRNA 2 *p* = 0.784 and *p* = 0.092 for 48 h and 72 h, respectively). (C) The distribution of cells in G1, S, and G2/M cell cycle phases in response to CRISPRi/a of *EPB41L4A-AS1* in HaCaT and A549 cells. Data are the difference in percentages of G1, S, and G2/M cells between cells transduced with target-specific gRNA 1 to those transduced with a control gRNA. Bars and error bars are mean and SEM of two or more independent replicates. ANOVA *p*-values were calculated from a hierarchical, linear model. HaCaT: G1: 1.7e-04, S: 0.052, and G2/M: 0.095; A549: G1: 0.011, S: 0.023, and G2/M: 0.59.

### *EPB41L4A-AS1* affects cell cycle phase distribution

In a previous study by our group, we reported that siRNA-mediated knockdown of *EPB41L4A-AS1* affected proliferation and cell cycle phase distribution (Hegre and Samdal *et.al.* unpublished). To investigate whether this effect was consistent between different gene perturbation methods, and not just an effect of the post-transcriptional downregulation, we used CRISPRi and CRISPRa to repress and activate the transcriptional level of *EPB41L4A-AS1,* respectively.

We investigated whether CRISPR-mediated modulation of *EPB41L4A-AS1* affected the viability of HaCaT cells by measuring the metabolic activity using an XTT viability assay. We observed a slight decrease in metabolic activity in HaCaT cells transduced with *EPB41L4A-AS1*-specific gRNAs compared to a non-specific gRNA control after 72 hours (Figure 2B), yet there were no significant effect between timepoints when we used CRISPRa to upregulate *EPB41L4A-AS1* (Supplementary Figure S2).

In line with previous results with siRNAs, we observed a decreased percentage of cells present in the G1 phase, and an increase in S and G2/M phases in response to CRISPRi-mediated downregulation of *EPB41L4A-AS1* in HaCaT cells (Figure 2C). As expected, the opposite was observed when *EPB41L4A-AS1* was upregulated, except for the G2/M phase, which displayed a slight enrichment of HaCaT cells in response to CRISPRa (Figure 2C). We used a hierarchical, linear model to calculate the combined effect on cell cycle phase distribution between CRISPRi and CRISPRa, requiring opposite contributions of the two assays and assuming a random effect for each assay. In HaCaT cells we observed a difference in cell cycle phase distribution for the G1 and S phases and these were consistent between the two assays. These changes were verified with another *EPB41L4A-AS1*-specific gRNA (Supplementary Figure S3). For the cell cycle assays we also included A549 cells to investigate whether the effect on cell cycle phase distribution was consistent across different cell lines. Compared with HaCaT cells, CRISPRi/a of *EPB41L4A-AS1* had an opposite effect in A549 cells, where downregulation of *EPB41L4A-AS1* led to a slight increase of cells present in the G1 phase, while the S and G2/M phases were not affected. Upregulation of *EPB41L4A-AS1* resulted in the opposite effect with a decreased percentage of A549 cells in the G1 phase. The number of cells in S phase was slightly increased, while the G2/M phase distribution was unaffected (Figure 2C).

### *EPB41L4A-AS1* regulates the expression of *EPB41L4A*

*EPB41L4A-AS1* is localized at chromosome 5 (hg38 chr5:112,160,526-112,164,818) and is transcribed from the opposite strand of *EPB41L4A,* where they partly overlap in a tail-to-tail orientation (Figure 3A). According to tissue expression data from the Genotype-Tissue Expression (GTEx) project, *EPB41L4A-AS1* is expressed across all tissues (Figure 3A; Supplementary Figure S4). Several antisense lncRNAs are known to regulate the expression of the protein coding gene that is overlapping on their opposite strand, and other neighboring genes, in *cis* [16]. To address whether *EPB41L4A-AS1* affects the expression of *EPB41L4A*, we used CRISPRi and CRISPRa-modified HaCaT cells transduced with an *EPB41L4A-AS1*-specific gRNA or a non-target gRNA. CRISPRi-mediated downregulation of *EPB41L4A-AS1* reduced the expression of *EPB41L4A* with two different gRNAs, whereas CRISPRa-mediated upregulation of *EPB41L4A-AS1* increased the expression of *EPB41L4A* (Figure 3B).

**Figure 3.**
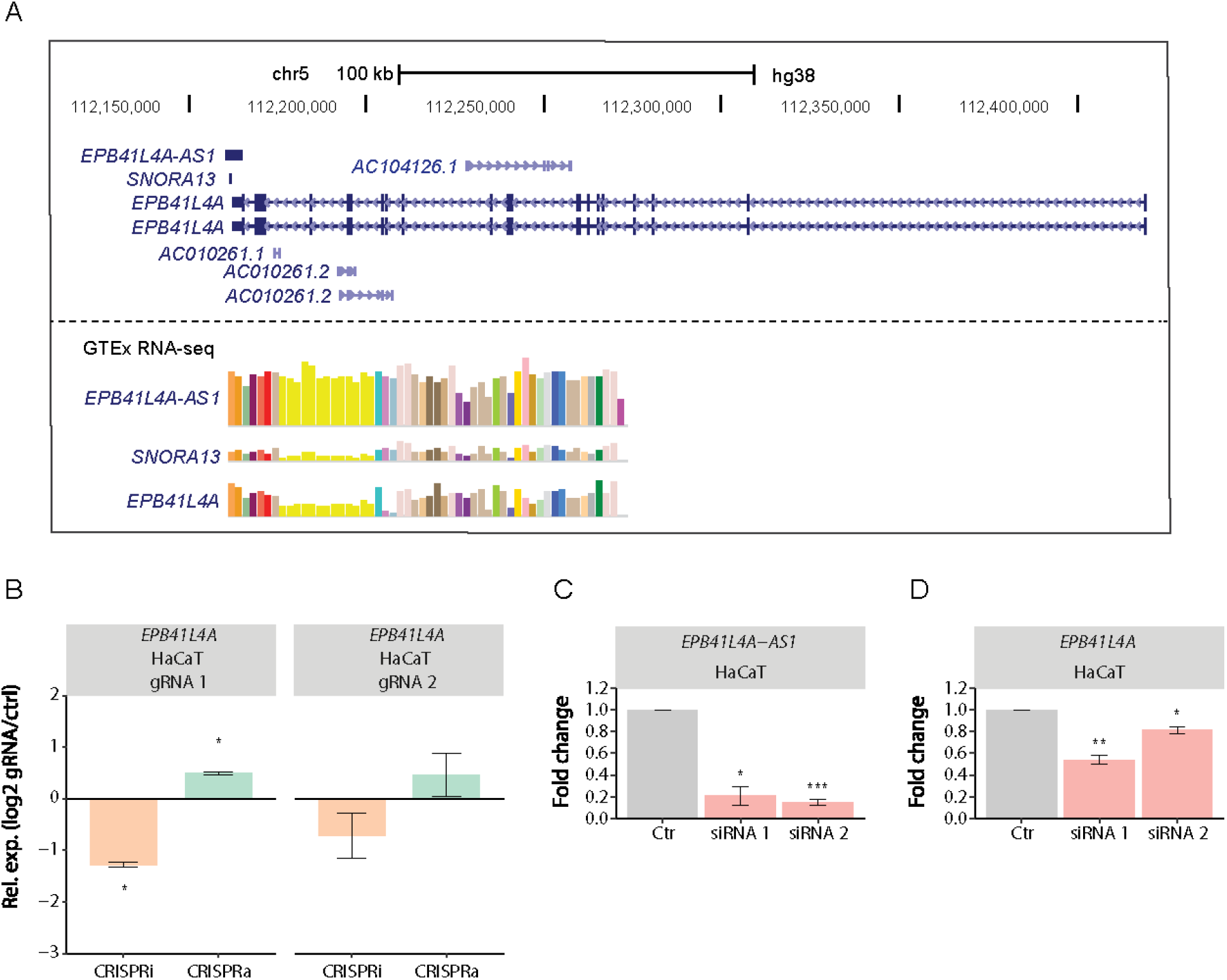
*EPB41L4A-AS1* is a positive regulator of *EPB41L4A*. (A) The genomic loci of *EPB41L4A-AS1* from the UCSC Genome Browser (https://genome.ucsc.edu/). Tissue expression data (transcript per kilobase million, TPM) are from the GTEx project. (B) The relative expression level of *EPB41L4A* as measured by RT-qPCR in response to CRISPRi/a of *EPB41L4A-AS1* in HaCaT cells. Data are presented as fold change expressions of cells transduced with target-specific gRNA 1 and gRNA 2 relative to control gRNA. Bars and error bars are mean and SEM of two independent replicates. Significant differences were determined by Student’s t-test, (unpaired, two-tailed) assuming equal variances (\**p* ≤ 0.05). (C) Percentage downregulation of *EPB41L4A-AS1* by using two siRNAs with different target sequences. Data are presented as fold change expressions of *EPB41L4A-AS1* following siRNA treatment relative to control-treated cells as measured by RT-qPCR in HaCaT cells. Bars and error bars are mean and SEM of three independent replicates. Significant differences were determined by Student’s *t*-test, (unpaired, two-tailed) assuming equal variances (\**p* ≤ 0.05; \*\*\**p* ≤ 0.001). (D) The relative expression level of *EPB41L4A* as measured by RT-qPCR in response to siRNA-mediated knockdown of *EPB41L4A-AS1* in HaCaT cells. Bars and error bars are mean and SEM of three independent replicates. Significant differences were determined by Student’s *t*-test, (unpaired, two-tailed) assuming equal variances (\**p* ≤ 0.05; \*\**p* ≤ 0.01).

We used two different *EPB41L4A-AS1*-specific siRNAs (Figure 3C) to investigate whether the effect of *EPB41L4A-AS1* on *EPB41L4A* was transcriptional or post-transcriptional. To disentangle nuclear from potential cytoplasmic effects, we designed these siRNAs such that siRNA 1 and siRNA 2 respectively targeted the first intron and the last exon of *EPB41L4A-AS1*; neither target site overlapped *EPB41L4A* exons. In line with the CRISPRi-mediated downregulation, both siRNAs knocked down *EPB41L4A-AS1* (Figure 3C) and caused a significant downregulation of *EPB41L4A* (Figure 3D). Combined, these results suggest that *EPB41L4A-AS1* positively regulates *EPB41L4A* at the post-transcriptional level within the nucleus.

### RNA sequencing identifies genes affected by CRISPRi of EPB41L4A-AS1

To get a mechanistic insight into how *EPB41L4A-AS1* affects viability and cell cycle phase distribution, we performed RNA-seq of CRISPRi-modified HaCaT and A549 cell lines transduced with *EPB41L4A-AS1*-specific gRNA or a non-target control gRNA (Figure 4A). Consistent with our RT-qPCR analyses, *EPB41L4A* was downregulated in both cell lines (*p*-value ≤ 0.05) in response to CRISPRi-mediated downregulation of *EPB41L4A-AS1* (Figure 4B). We identified a total of 1612 and 1371 differentially expressed genes with *p*-value ≤ 0.05, where 8 and 13 of these genes had an adjusted *p*-value ≤ 0.05, in HaCaT and A549 cells, respectively (Figure 4C). Moreover, in the joint analysis of HaCaT and A549 cells we identified 1439 differentially expressed genes with *p*-value ≤ 0.05 (Figure 4C; Supplementary Dataset S1). Gene ontology (GO) enrichment analyses of the differentially expressed genes in HaCaT, A549, or the joint analysis of CRISPRi vs. control showed enrichment of signaling pathways (KEGG and Reactome) associated with proliferation, including NTRK, ERK, and ARMS for the upregulated genes namely *TMEM189-UBE2V1*, *KIDINS220*, *C1orf167-AS1*, *ZSWIM8-AS1*, and *RMRP* (Figure 4D).

**Figure 4.**
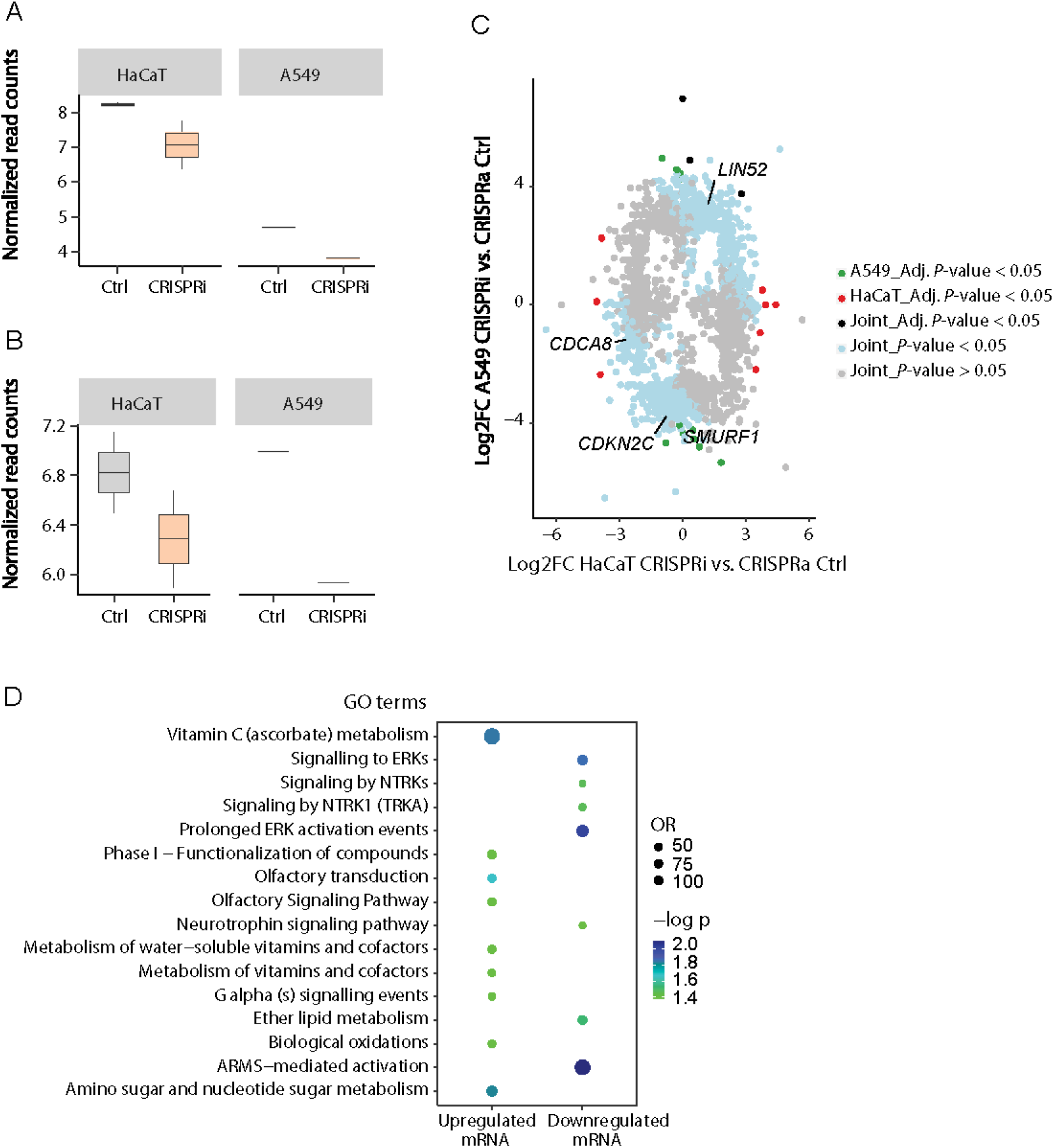
*EPB41L4A-AS1* affects gene expression. (A) Box plot showing RNA-seq expression levels of *EPB41L4A-AS1* in CRISPRi-modified HaCaT (n=2) and A549 cells (n=1) transduced with an *EPB41L4A-AS1*-specific gRNA compared to a non-specific gRNA control. Data are presented as normalized read counts. Median is indicated by the central line in the boxes and whiskers indicate the first and third quartiles (25th and 75th percentiles). (B) Box plot showing RNA-seq expression levels of *EPB41L4A* in CRISPRi-modified HaCaT (n=2) and A549 cells (n=1) transduced with an *EPB41L4A-AS1*-specific gRNA compared to a non-specific gRNA control. Data are presented as normalized read counts. Median is indicated by the central line in the boxes and whiskers indicate the first and third quartiles (25th and 75th percentiles). (C) Differentially expressed genes in CRISPRi compared to control in the individual analysis of HaCaT and A549 cells. There were 2871 differentially expressed genes colored according to *p*-values in individual and joint analysis (CRISPRi vs. Control in the joint data from HaCaT and A549 cell lines). (D) GO enrichment analysis of differentially expressed genes (adjusted *p*-value < 0.05) in either A549, or HaCaT, or in the joint analysis, combined by requiring the same effect direction in each comparison. The terms are from KEGG and Reactome. The colors represent the negative log10 of adjusted *p*-values, so dark blue color represents more significant GO terms. Size of the dots represent the odds ratio (OR).

## Discussion

Our data show that *EPB41L4A-AS1* is localized as bright foci both in and outside the nucleus in HaCaT cells. Consistent with the nuclear foci having a role in gene regulation, we found that *EPB41L4A-AS1* positively regulates the expression of its antisense mRNA *EPB41L4A*, as CRISPR-mediated up- and downregulation of *EPB41L4A-AS1*, up- and downregulates *EPB41L4A*, respectively. Moreover, this effect is post-transcriptional, as we found that siRNA-mediated knockdown of *EPB41L4A-AS1* also downregulates *EPB41L4A*.

In line with our previous study, CRISPR-mediated downregulation of *EPB41L4A-AS1* expression affected cell cycle phase distribution with a reduction of HaCaT cells in the G1 phase, while CRISPRa-mediated upregulation led to an increased percentage of cells in the G1 phase. The effect on cell cycle phase distribution is cell type-specific, as CRISPR-mediated decrease and increase of *EPB41L4A-AS1* expression in A549 cells resulted in an increase and decrease of cells in G1 phase, respectively. Finally, we found that downregulation of *EPB41L4A-AS1* affects the gene expression of multiple genes in both HaCaT and A549 cells, indicating that *EPB41L4A-AS1* has both *cis*- and *trans* regulatory abilities. Several of the affected genes are involved in signaling pathways associated with proliferation, possibly explaining *EPB41L4A-AS1*’s proliferation phenotype.

The subcellular localization of *EPB41L4A-AS1* was both nuclear and extranuclear in HaCaT cells. The extranuclear localization appears to be cytoplasmic, which is in line with *EPB41L4A-AS1* encoding a small protein, TIGA1. TIGA1 is translated from the first exon of *EPB41L4A-AS1,* and a study demonstrated that ectopic expression of TIGA1 reduced colony-formation and growth of the lung cancer cell line EBC-1 in soft agar [22]. Whereas the extranuclear foci suggests that TIGA1 also is present in HaCaT cells, our previous (Hegre and Samdal et al., unpublished) and current results indicate that downregulating *EPB41L4A-AS1* negatively affects HaCaT proliferation. Consequently, either TIGA1 has cell type-dependent effects on proliferation or *EPB41L4A-AS1* affects proliferation through other mechanisms beside TIGA1.

A likely such mechanism is *EPB41L4A-AS1* directly regulating nuclear gene expression, as we show that *EPB41L4A-AS1* localizes to nuclear DNA and that *EPB41L4A-AS1* positively regulates the expression of its antisense mRNA *EPB41L4A.* EPB41L4A is a part of a membrane-associated protein superfamily that contains a FERM (Four-point-one, Ezrin, Radixin, Moesin) domain. These proteins are important for embryonic development and have regulatory functions involving intracellular trafficking, signal transductions, and cytoskeletal rearrangement [23]. Moreover, *EPB41L4A* is a target gene of the Wnt/β-catenin pathway [24], a pathway that regulates embryonic development and adult tissue homeostasis, in addition to proliferation and differentiation [25]. Depending on their genomic configuration, sense-antisense genes can share regulatory DNA elements that jointly control their gene expression. However, for *EPB41L4A-AS1* and *EPB41L4A* the distance between their transcription start sites (TSS) is 258751 bb (Figure 3A). Although the two TSSs could still be close in three-dimensional space in the nucleus, our result showing that siRNA-mediated knockdown of *EPB41L4A-AS1* also downregulates *EPB41L4A* strongly supports that the regulation is post-transcriptional and caused by the RNA of *EPB41L4A-AS1*. Indeed, *EPB41L4A-AS1*’s bright foci that co-localize with the DNA in HaCaT cells are consistent with an accumulation of *EPB41L4A-AS1* at its site of transcription.

As mentioned above, we found that CRISPR-mediated downregulation of *EPB41L4A-AS1* slightly reduced the viability in HaCaT cells, which supports results from our recent study where siRNA-mediated knockdown of *EPB41L4A-AS1* reduced proliferation of the colorectal cancer (CRC) cell line DLD1, as well as in HaCaT and A549 cells (Hegre and Samdal et.al. unpublished). In line with our results, Bin et. al. found that knockdown of *EPB41L4A-AS1* in the CRC cell lines HCT116 and SW620 reduced proliferation, while overexpression of *EPB41L4A-AS1* had the opposite effect. Moreover, the expression of *EPB41L4A-AS1* was higher in CRC tissue compared to normal tissue. They concluded that *EPB41L4A-AS1*functions as an oncogene by regulating the Rho/ROCK pathway [26]. Another study identified *EPB41L4A-AS1* as a possible hub lncRNA in colorectal cancer [9], indicating a potential role as a prognostic marker.

We further validated that *EPB41L4A-AS1* is involved in the regulation of cell cycle progression by using CRISPRi to downregulate *EPB41L4A-AS1* in HaCaT and A549 cells. In addition, we strengthened the evidence by using CRISPRa to upregulate *EPB41L4A-AS1,* which affected the cell cycle phase distribution in the opposite direction. Moreover, the effect on cell cycle phase distribution was different in HaCaT and A549 cells, suggesting cell type-specific functions of *EPB41L4A-AS1.* Other recent studies indicate that the expression of *EPB41L4A-AS1* and its biological role seems to vary in different types of cells, tissues, and cancer stages. A study from Rao et.al. found a correlation between low expression of *EPB41L4A-AS1* and early-stage breast cancer [8], whereas two other studies reported an upregulation of *EPB41L4A-AS1* in cervical and ovarian cancer [27, 28]. One of the studies identified *EPB41L4A-AS1* as a central hub in a sub-network in cervical cancer. Analysis indicated an interaction between *EPB41L4A-AS1* and the genes *CCND2* and *VDAC1* via different microRNAs, proposing a potential role as a biomarker for disease progression and as a therapeutic target [27]. In the study from Zhou et.al, *EPB41L4A-AS1* was categorized as a risky lncRNA linked to stage progression and poor clinical outcome in ovarian cancer, and might be a potential candidate for classification of patients into subgroups [28].

In some functional studies, *EPB41L4A-AS1* was reported to be involved in metabolic reprogramming and as a repressor of the Warburg effect in placental tissue of miscarriage [12] and in cancer cells (cervical, breast, bladder and liver) [11]. Liao et.al. found that *EPB41L4A-AS1* was regulated by p53 at the transcriptional level by direct binding to the promoter in Hela and HepG2 cells. Also, stable knockdown of *EPB41L4A-AS1* in HeLa cells increased tumor colony formation, a process that was dependent on the presence of glutamine metabolism. Furthermore, an increased therapeutic effect of a glutaminase inhibitor in *EPB41L4A-AS1* knockdown cells was confirmed in vivo in mice [11]. Another study identified *EPB41L4A-AS1* as a direct *MYC* target gene, which was downregulated in response to MYC overexpression in P493-6 human B-cells [29].

As *EPB41L4A-AS1* has been detected as dysregulated in several cancers, with functions affecting several cellular processes, including metabolic reprogramming, tumor colony formation, gene regulation, cell cycle, and proliferation, it probably has *trans* regulatory abilities. We found that downregulation of *EPB41L4A-AS1* resulted in differentially expressed genes in both A549 and HaCaT cells, including several genes involved in proliferation, such as *SMURF1, F2RL1, CPNE3,* and *KIDINS220*. Moreover, GO terms included several growth-promoting pathways, including NTRK, ERK, and ARMS mediated signaling.

In summary, *EPB41L4A-AS1* is located both in the nucleus and cytoplasm in HaCaT cells. We demonstrate that *EPB41L4A-AS1* has a slight effect on cellular metabolic activity and that it exhibits cell type-specific functions as it affects the cell cycle phase distribution differently in HaCaT and A549 cells. Finally, *EPB41L4A-AS1* regulates the expression of its antisense mRNA *EPB41L4A* through post-transcriptional effects in the nucleus and downregulating *EPB41L4A-AS1* affects more than a thousand other genes in HaCaT and A549 cells, including genes involved in proliferation and growth-promoting signaling pathways.

## Acknowledgement

This work was supported by the Norwegian Cancer Society (grant number 2278701) and the Research Council of Norway (grant number 230338). The library preparation and sequencing were provided by the Genomics Core Facility (GCF), Norwegian University of Science and Technology (NTNU). GCF is funded by the Faculty of Medicine and Health Sciences at NTNU and Central Norway Regional Health Authority. The light microscopy was provided by the Cellular and Molecular Imaging Core Facility (CMIC), NTNU. CMIC is funded by the Faculty of Medicine at NTNU and Central Norway Regional Health Authority.

## Author contribution

HS planned and performed imaging and wet lab experiments, contributed with statistical analysis, and wrote the manuscript. SAH contributed with statistical analysis, generated all the figures and was a major contributor in editing the manuscript. KC analyzed the RNA-seq data. NBL performed all FACS analysis. PAA contributed with the planning and design of CRISPR experiments. BS contributed to the planning of RNA FISH imaging and performed the image analysis. PS contributed to the study design, supervised the project, and edited the manuscript. All authors read and approved the final manuscript.

## Supplementary Data

**Supplementary file 1:** Supplementary Figures and Tables

**Supplementary dataset 1**: RNA-seq identified a total of 1439 differentially expressed genes with *p*-value ≤ 0.05 in the joint analysis of HaCaT and A549 cells.

