## Supplementary file 1 for "The lncRNA *EPB41L4A-AS1* regulates gene expression in the nucleus and exerts cell type-dependent effects on cell cycle progression"

### Supplementary Figures

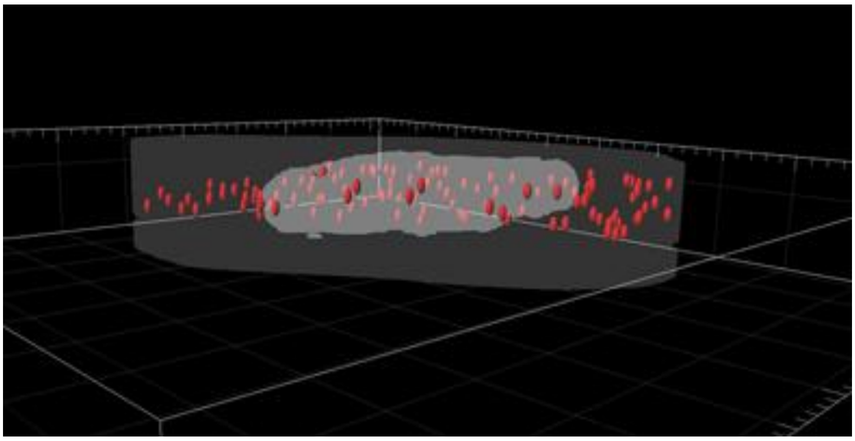

**Supplementary Figure S1: Cross-sections of the nucleus in HaCaT cells show *EPB41L4A-AS1* (red) localized with DNA (DAPI, grey).**

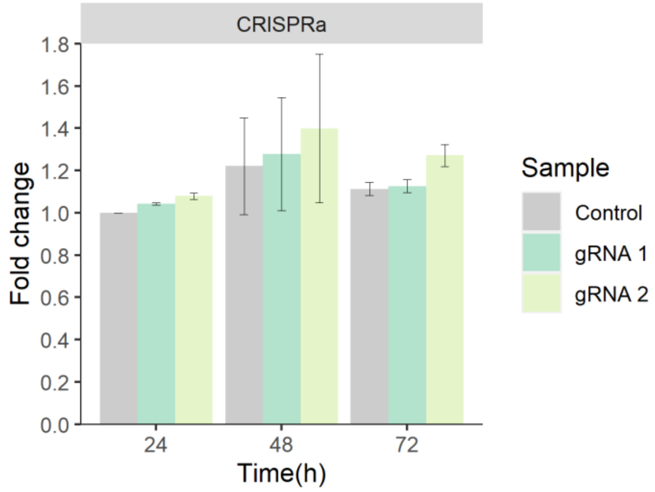


**Supplementary Figure S2: Effect of CRISPRa of *EPB41L4A-AS1* on the metabolic activity in HaCaT cells, as measured by XTT.** Data are presented as normalized absorbance ($A_{465\ nm} - A_{630\ nm}$). Bars and error bars are mean and standard error of mean (SEM) of three independent replicates.

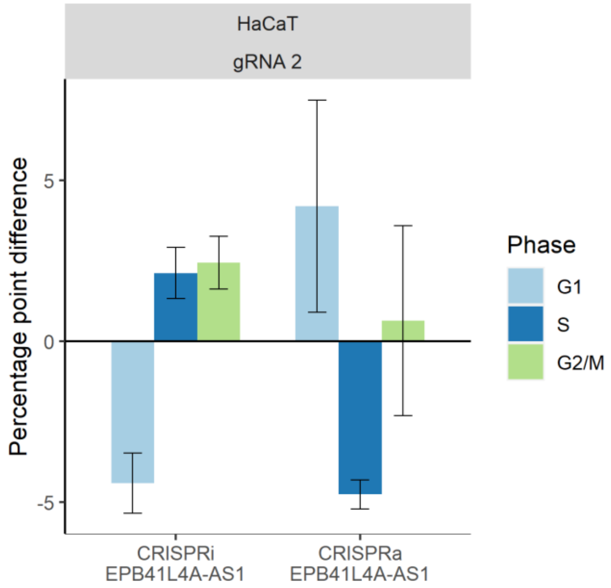


**Supplementary Figure S3: Distribution of HaCaT cells in G1, S, and G2/M cell cycle phases in response to CRISPRi/a of *EPB41L4A-AS1*.** Data are the difference in percentages of G1, S, and G2/M cells between cells transduced with target-specific gRNA 2 to those transduced with a control gRNA. Bars and error bars are mean and SEM of two or more independent replicates. ANOVA *p*-values were calculated from a hierarchical, linear model: G1: 6.2e-04, S: 0.009, and G2/M: 0.173.

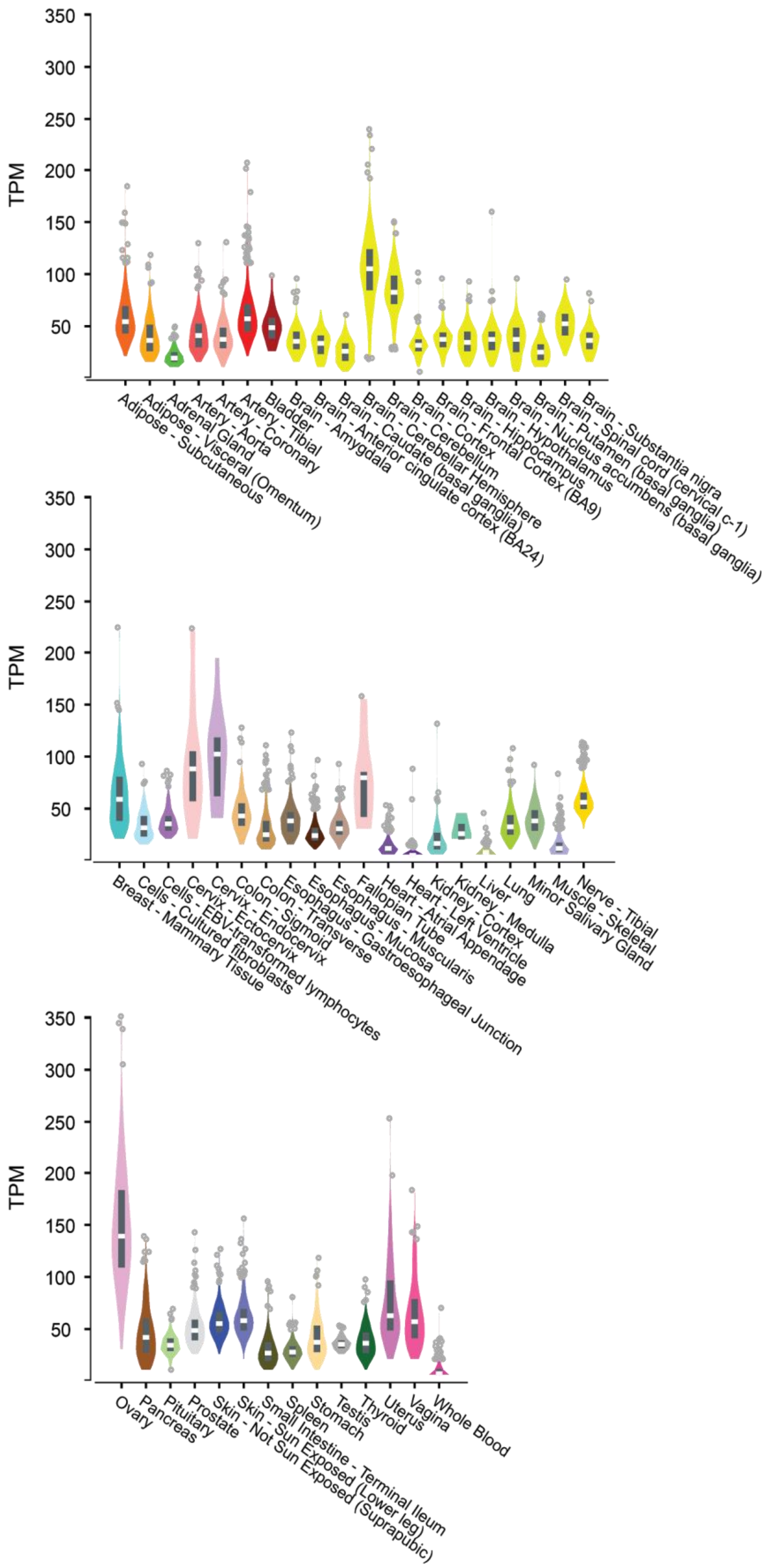


**Supplementary Figure S4: Tissue expression (transcript per kilobase million, TPM) for *EPB41L4A-AS1* (ENSG00000224032.6) from the Genotype-Tissue Expression (GTEx) project.**

#### Supplementary Tables

##### Supplementary Table S1: Custom Stellaris FISH probes conjugated to a Quasar670 dye in the 3′ end.

| Probe sequences |
|---|
| cacgtattttgggacccgac |
| aggagcccaaaggactttag |
| gagcatatgatcagtgctgg |
| agaggaggtatccagacatc |
| gatgtgtggccagttctttc |
| gtcaccaaagggaggaggag |
| gagggagaagtgccagtgag |
| ttctacaagtggaaggaccc |
| aagaaaggacaggcttccgt |
| acagcaagctcgcaccaaaa |
| cagaggacccagagcaaaaa |
| agtcgactcaccagactcag |
| ttccaaacgtgggaacgcag |
| ctggcatagtcgatgatgta |
| aaccaggcttatctggagaa |
| aggccttttcactgacgaaa |
| gggcaagcataaagtcagtt |
| gcacgctgcttttaagttat |
| gcacgtggaaacagctgtaa |
| tgtagggtaggtgatcactc |
| ctacacttcaggcatccatc |
| agatggtattcagctagcag |
| ttcaggtcacctttatgctc |
| caaaatcctcaccaaggaca |
| ggcaactcttgtgaagatca |
| tcagactcccagaaatttca |
| caggtaattatgtcagtact |
| aatgtgtatttacagactcc |
| ttattcacatcctcactgtc |
| acacaaatgccaaagtgcat |
| gtttctggcagttaatcact |
| tcctcccttataacttttta |
| ggtgacagcagtgaatactg |
| acacaagctattttaaggct |
| aagtacatttctctttgggc |
| tagccacgatttttgagtca |

**Supplementary Table S2: Oligos used for guide RNA (gRNA) cloning.**

gRNAs were ordered from Sigma-Aldrich

| Gene primer | Oligo sequences (gRNA sequences in bold) | CRISPRi/a |
|---|---|---|
| neg_ctr_F | CACCG**TGCGATGGGGGGGTGGGTAGC** | Negative control (ctr) |
| neg_ctr_R | AAAC**GCTACCCACCCCCCCATCGCA**C | |
| MALAT1 _F | CACCG**CAGCCCGAGACTTCTGTAA** | Positive ctr CRISPRi |
| MALAT1 _R | AAAC**TTACAGAAGTCTCGGGCTG**C | |
| SLC4A1 _F | CACCG**TCAGGAGAACCATGGGGACC** | Positive ctr CRISPRa |
| SLC4A1_R | AAAC**GGTCCCCATGGTTCTCCTGA**C | |
| Epb_AS1_1_F (gRNA1) | CACCG**GCGTCTCGCCATAGCGCAG** | CRISPRi/a |
| Epb_AS1_1_R | AAAC**CTGCGCTATGGCGAGACGC**C | |
| Epb_AS1_2_F (gRNA2) | CACCG**TGACTCGGGCTGAGAACCC** | CRISPRi/a |
| Epb_AS1_2_R | AAAC**GGGTTCTCAGCCCGAGTCA**C | |

F = forward, R = reverse

**Supplementary Table S3: siRNAs used in RNA interference experiments.**

| Target lncRNA | Alias | Cat. No. | Producer | Sense sequence |
|---|---|---|---|---|
| SiRNA neg control | Ctr | SIC001 | Sigma | |
| EPB41L4A-AS1 | E1 | CTM-431274 | Dharmacon | ACAGUGAGGAUGUGAAUAAUU |
| EPB41L4A-AS1 | E2 | CTM-431275 | Dharmacon | GUGAGGAUGUGAAUAAAUAUU |

**Supplementary Table S4: Primers used for RT-qPCR.**

Qiagen QuantiTect Primer Assays (249900) were used for mRNA expression analysis and Qiagen $RT^2$ lncRNA PCR assays (330701) were used for lncRNA expression analysis.

| Gene | Primer assay | Cat.no |
|---|---|---|
| GAPDH | Hs_GAPDH_1_SG QuantiTect Primer Assay | QT00079247 |
| EPB41L4A | Hs_EPB41L4A_1_SG QuantiTect Primer Assay | QT00101003 |
| SLC4A1 | Hs_SLC4A1_1_SG QuantiTect Primer Assay | QT00068502 |
| MALAT1 | $RT^2$ lncRNA qPCR Assay for Human MALAT1 | LPH18065A |
| EPB41L4A-AS1 | $RT^2$ lncRNA qPCR Assay for Human EPB41L4A-AS1 | LPH04387A |
